# Global transcriptional analysis of *Geobacter sulfurreducens* under palladium reducing conditions reveals new key cytochromes involved

**DOI:** 10.1101/319194

**Authors:** Alberto Hernández-Eligio, Aurora M. Pat-Espadas, Leticia Vega-Alvarado, Manuel Huerta-Amparán, Francisco J. Cervantes, Katy Juárez

## Abstract

*Geobacter sulfurreducens* is capable of reducing Pd(II) to Pd(0) using acetate as electron donor; however, the biochemical and genetic mechanisms involved in this process have not been described. In this work, we carried out transcriptome profiling analysis to identify the genes involved in Pd(II) reduction in this bacterium. Our results showed that 252 genes were upregulated while 141 were downregulated during Pd(II) reduction. Among the upregulated genes, 12 were related to energy metabolism and electron transport; 50 were classified as involved in protein synthesis; 42 were associated with regulatory functions and transcription, and 47 have no homologs with known function. RT-qPCR data confirmed upregulation of genes encoding PilA, the structural protein for the electrically conductive pili, as well as *c*-type cytochromes GSU1062, GSU2513, GSU2808, GSU2934, GSU3107, OmcH, OmcM, PpcA, PpcD under Pd(II)-reducing conditions. Δ*pilA* and Δ*pilR* mutant strains showed 20% and 40% decrease in the Pd(II)-reducing capacity, respectively, compared to the wild type strain, indicating the central role of the pili in this process. RT-qPCR data collected during Pd(II) reduction also confirmed downregulation of the *omcB, omcC, omcZ,* and *omcS* genes, which have been shown to be involved in Fe(III) reduction and electrodes. Based on these results, we propose mechanisms involved in Pd(II) reduction by *G. sulfurreducens*.

**Importance:** *Geobacter sulfurreducens* is a versatile microorganism, known for its ability to reduce a wide range of environmentally relevant metals. It has been reported that this bacterium synthesizes palladium nanoparticles successfully. Yet, the biochemical and genetic mechanisms involved in this process had not been described previously. By using a transcriptome profile analysis to identify genes implicated in this process and genetic and physiological data, we propose a model for the biological reduction of Pd(II) by *G. sulfurreducens*. Moreover, the study also revealed the microbial reduction of Pd(II) coupled to growth, which shows not only unexpected ideas about its complex metabolism, but some key cytochromes involved in this pathway, which have not been previously associated with other metal reducing process for this bacterium. In brief the present work contributes to a better understanding of the biochemical and genetic mechanisms involved in Pd(II) reduction and put forward novel insights about *G. sulfurreducens* metabolism.

## Introduction

In the last two decades, the alternative of using bioreductive deposition of precious metals, such as the platinum-group (PGMs; e.g Pt, Pd, Rh), for their recovery has been widely explored. Special interest has been focused on palladium (Pd) due to its high value and extensive use as catalyst. Fe(III) reducing bacteria (IRB), such as *Shewanella oneidensis* (1) and *Geobacter sulfurreducens* (2), as well as sulfate reducing bacteria (SRB), such as *Desulfovibrio desulfuricans* (3), have extensively been explored for this purpose. Despite the great demand for developing efficient microbial processes to recover this valued element, the mechanisms involved in Pd(II) reduction and subsequent deposition of Pd(0) are poorly understood.

Extracellular formation of Pd(0) nanoparticles (NP’s) has been reported in *G. sulfurreducens* (2). Moreover, differences regarding location of deposited NP’s, depending on the strain, have been documented (3). These findings could be related to specific mechanisms used by each strain, as well as to experimental conditions prevailing; however, further studies are required to fully elucidate the mechanisms involved.

*Geobacter* species are abundant in nature and have the ability to perform extracellular electron transfer (EET) to reduce a broad array of heavy metals, such as Fe(III), Mn(IV), U(VI), Co(III) and Ag(I), among others (4, 5, 6, 7). Regarding Pd(II) reduction, using acetate as electron donor, *G. sulfurreducens* has been used to reduce up to 25 mg Pd(II)/L with the concomitant deposition of Pd(0) NP’s (2). It has been proposed that the electron transfer reported for iron reduction could also be involved in Pd(II) reduction (8).

Nevertheless, the genome of *G. sulfurreducens* has 111 predicted c-type cytochromes (9, 10), hence it is likely that other cytochromes, as well as conductive pili, could play a role in Pd(II) reduction (11, 12). This is supported by the observation that some proteins or cytochromes are specifically required to achieve the reduction of certain metal species, as it has been reported for soluble Fe(III), Fe(III) oxides and U(VI) (9, 13, 14).

The purpose of this study was to elucidate the mechanisms involved in Pd(II) reduction by *G. sulfurreducens*, based on global transcriptome analysis using RNA sequencing analysis. Quantitative reverse transcription PCR, genetic and physiological tests were also performed to identify genes involved in Pd(II) reduction. The Pd(II) reduction pathways in *G. sulfurreducens* were proposed based on the obtained results.

## Materials and Methods

### Culture procedures

Bacterial strains and oligonucleotides used in this study are listed in Table 1. *G. sulfurreducens* PCA was grown anaerobically at 30 °C in NBAF medium, supplemented with acetate and fumarate; these culture conditions were referred to as “Non-Pd(II)-reducing conditions” (15). For experiments conducted under “Pd(II)-reducing” conditions, late logarithmic phase cultures of *G. sulfurreducens* were used. The protocol consisted of harvesting cells by centrifugation at 9,000 *g* for 20 min and washing with a sterilized, osmotically balanced buffer. The reaction buffer composition was as follows (in grams per liter): NaHCO_3_, 2.5; NH_4_Cl, 0.25; NaH_2_PO_4_·H_2_O, 0.006; and KCl, 0.1. Reduction experiments were performed in 120-ml glass serum bottles capped with rubber stoppers including 100 ml of anaerobic basal medium, flushed with N_2_/CO_2_ (80:20, v/v). Cell suspensions, as well as acetate and Na_2_PdC_4_ (Sigma-Aldrich) stock solutions were added to yield a concentration of 800 mg l^-1^ cell dry weight (CDW), 5 mM and 25 mg l^-1^, respectively (2, 8). RNA Later (Ambion) was added to cultures and cells for harvesting cells. Bacterial pellets were flash frozen and stored at - 70 °C.

**TABLE 1.**
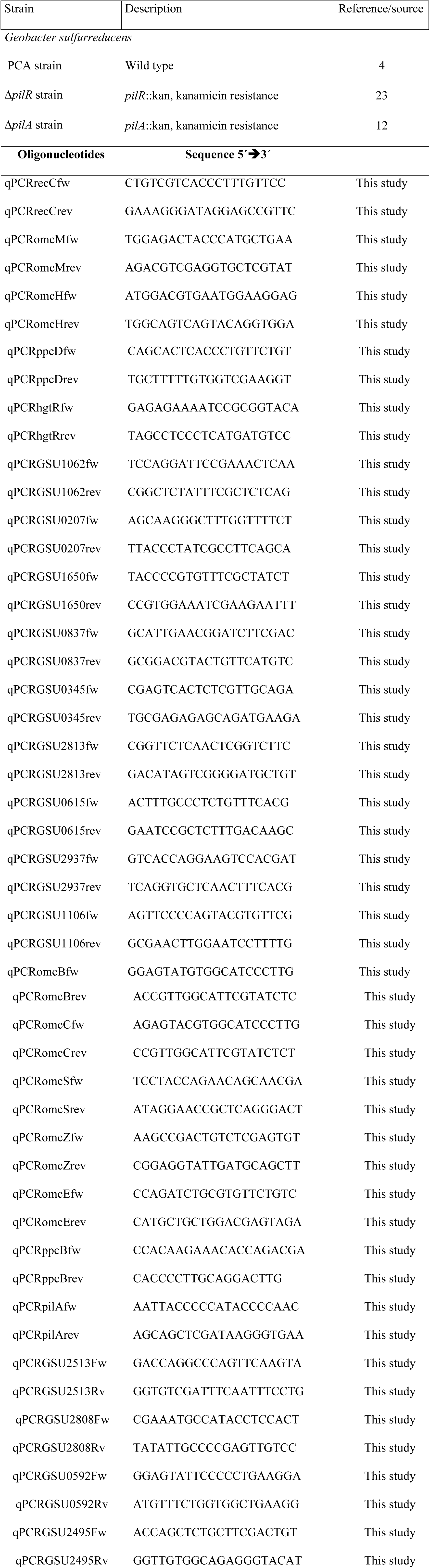
Bacterial strain and primer sequences used in this work and for RT-qPCR validations

### RNA extraction

*G. sulfurreducens* cells from both “Pd(II) reduction” and “Non-Pd(II) reduction” conditions were used for RNA-Seq and RT-qPCR analyses. All experiments were performed in duplicates. For each biological sample, total RNA samples were extracted using the RNeasy mini kit (Qiagen), then they were examined with an Agilent 2100 Bioanalyzer and quantified using NanoDrop 200c (Thermo Scientific).

### RNA-Seq and data analysis

RNA-Seq was performed using RNA samples extracted from two independently grown cultures with and without Pd(II). Illumina sequencing was performed at the USMI (Unidad de Secuenciación Masiva, UNAM, Mexico). Briefly, after removing residual DNA using DNase I (ThermoScientific) and ribosomal RNA with Terminator 5′-Phosphate-dependent exonuclease (Epicentre), the mRNA-enriched RNA was chemically fragmented to 150~200 bp. Based on these cleaved RNA fragments, cDNA was synthesized using a random hexamer primer and reverse transcriptase. After end reparation and ligation of adaptors, the products were amplified by PCR, further purified, and used to create the final cDNA library. Libraries were sequenced on an Illumina Genome Analyzer IIx. Differential expression analyses were performed through IDEAmex website (http://zazil.ibt.unam.mx/ideamex/) using three methods: edgeR (16), DESeq (17) and NOISeq (18). edgeR and NOISeq were performed by applying TMM (19) as the normalization method. To identify differentially expressed genes, we selected those whose p value was <0.05 and fold change > 2, for each method. Finally, we considered as the best candidates, only genes that appeared differentially expressed in the three methods. The functional annotation of differentially expressed genes regarding the affected pathways was obtained from Kyoto Encyclopedia of Genes and Genomes (KEGG) (20), using our own R’s scripts. RNA-Seq transcriptome data were deposited in the NCBI Gene Expression Omnibus database under accession number GSE113152.

### Quantitative real-time RT-PCR (RT-qPCR)

To validate the quality of the sequencing data, some differentially expressed genes were selected for RT-qPCR analysis. mRNA was extracted as described in the section “RNA extraction” and DNA residual was removed using DNase I (Thermo Scientific). cDNA synthesis was performed using RevertAid H Minus First Strand cDNA Synthesis kit (Thermo Scientific). Subsequently, the RT-qPCR was performed using a Maxima SYBR Green/ROXq PCR Master Mix (Thermo Scientific) in a 96-well plate with the Light-Cycler II (Roche). Gene-specific primers used for RT-qPCR are shown in Table 1. *recC* was used as gene internal standard for PCR amplification. Normalized fold changes of the relative expression ratio were quantified by the 2^-ΔΔCT^ method (21). All experiments were performed in triplicates and their average values were calculated.

### Cytochrome c content

Membrane fractions of *G. sulfurreducens* were isolated as previously described (22, 23). Outer membrane-enriched fractions were prepared by treating crude membranes with a sarkosyl (sodium N-lauroyl sarcosinate solution at 1% wt/vol) to extract inner membrane proteins. Outer membrane proteins were analyzed by Tris-Glycine denaturing polyacrylamide gel electrophoresis, and *c*-type cytochromes were detected by staining with N,N,N,N-tetramethylbenzidine, as previously described (24, 25). PageRuler pre-stained protein standards were purchased from Thermo Scientific. The Tris-Glycine gel image was digitized using a Gel-doc (Bio Rad).

### Immunoblot analysis

Protein extraction from cultures performed under “Pd(II)-reducing” and “Non-Pd(II)-reducing” conditions was conducted by western blot as follows: cells pellets were resuspended in 150 μl of B-PER II Bacterial Protein extraction reagent (Pierce) and incubated for 15 min. 1.0 mg of total protein per sample was incubated with PAGE-Buffer and boiled for 5 min and were separated on a 15% SDS-PAGE. After separation, proteins were transferred to nitrocellulose membranes (Merck-Millipore) for immunoblot analysis using rabbit polyclonal antibodies raised against *G. sulfurreducens* PilA. Blots were blocked with 3% BSA in PBS overnight at 4°C and then incubated with a 1/1,000 dilution of primary antibody for 4 h at room temperature, washed with PBS, and incubated with a 1/5,000 dilution of goat anti-rabbit alkaline phosphatase-conjugated secondary antibody for 3 h at room temperature. After being washed, blots were developed with 5-bromo-4-chloro-3- indolylphosphatase (BCIP)-Nitro Blue Tetrazolium (Pierce) following the manufacturer’s instructions.

### Viability assay

Cell viability assays after exposure to Pd(II) was performed by recovering the resting cells in NBAF medium and incubated to measure microbial growth. Prior to inoculation, cell suspensions under Pd(II)-reducing conditions were incubated for 3 h and further transferred from there to NBAF medium with a cellular density of 0.05 (OD 600_nm_). The cultures were incubated at 30 °C, and growth was periodically monitored as OD 600_nm_.

### Microbial growth under Pd(II)-reducing conditions

*G. sulfurreducens* was grown in bicarbonate-buffered medium (26), supplemented with 15 mM pyruvate, 5 mg l^-1^ of Na_2_PdCl_4_ and 15 mM glutamine. Cultures were incubated at 30 °C, and Pd(II) reduction, pyruvate consumption and growth were periodically monitored. All experimental treatments were set-up in triplicate.

### Analytical techniques

Reduction of Pd(II) was quantified as follows: 5 ml of Pd(II) medium, inoculated or non-inoculated, were filtered using 0.22 μm membrane filters (Millipore, Bedford, USA).

Filtrate samples were then analyzed by inductively coupled plasma-optic emission spectroscopy (ICP-OES, Varian 730-ES).

Pyruvate consumption was measured by a high performance liquid chromatography (HPLC, Agilent Technologies 1200 Series, Santa Clara, CA, USA) equipped with an Aminex HPX-87H column (BIO-RAD). The column was maintained at 50°C and was eluted with 5 mM H_2_SO_4_ at a flow rate of 0.6 ml min^-1^. Pyruvate was detected by an UV detector at 210 nm. The standard reagent for quantification was purchased from Sigma-Aldrich (St Louis, MO, USA).

### X-ray diffraction analysis

Analysis of the Pd(0) NP’s deposited on the different strains of *G. sulfurreducens* was conducted in an X-Ray diffractometer Bruker D8 Advance. Samples were treated as previously described (2). X-ray diffraction (XRD) patterns were recorded from 20°-90° 2**θ** with a step time of 2 s and step size of 0.01° 2**θ**.

## Results and Discussion

### Differentially expressed genes during Pd(II) reduction

Previous studies have shown that *G. sulfurreducens* is able to reduce Pd(II) to Pd(0) NP’s (2). However, the proteins involved in this respiratory process have not been identified. To examine the transcriptional changes that occur in *G. sulfurreducens* during Pd(II) reduction, we used RNA-Seq. The p-value and fold change were calculated by the statistical methods DESeq, edgeR and NOISeq. Only genes that showed differential expression by all methods were selected, resulting in 393 differentially expressed genes (Fig. 1A). A cut off p-value <0.05 and a fold change >2 was used.

**FIG 1.**
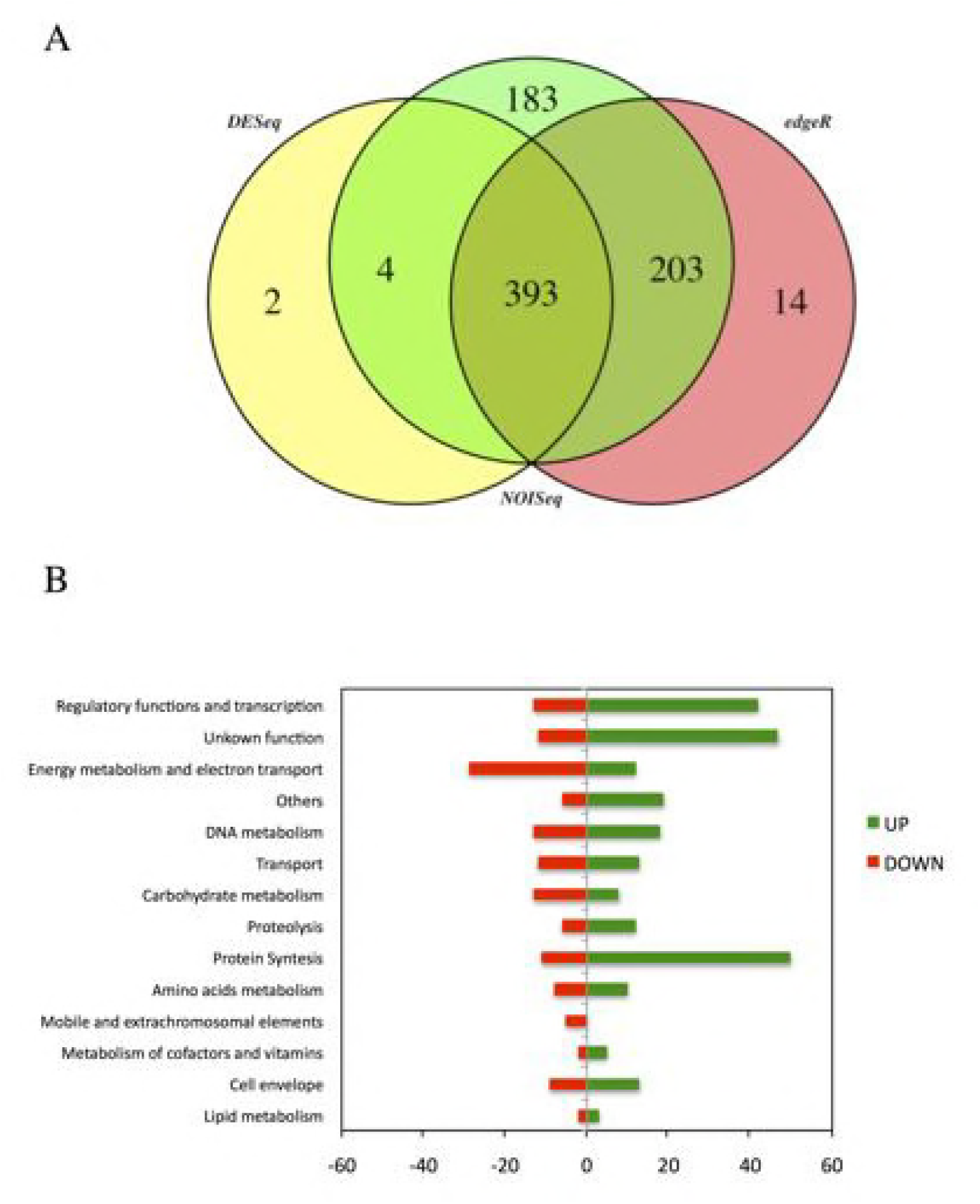
Transcriptome analysis results from *Geobacter sulfurreducens* under Pd(II)-reducing conditions. (A) Venn diagram representing differential gene expression analysis from Pd(II)-reducing conditions by three statistical methods. (B) Functional overview of the genes that were differentially expressed during Palladium reduction.

Out of the 393 genes, 252 displayed statistically significant upregulation (>2 FC), and 141 downregulation under Pd(II)-reducing conditions. This accounted for approximately 11% of the genes in the *G. sulfurreducens* genome, indicating that Pd(II) reduction triggered significant global gene expression changes. The genes that showed significant differences in transcript levels were classified into the following functional categories: regulatory functions and transcription; energy metabolism and electron transport; DNA metabolism; transport; carbohydrate metabolism; proteolysis; protein synthesis; amino acids metabolism; mobile and extrachromosomal elements; metabolism of cofactors and vitamins; cell envelope; lipid metabolism; unknown function; and others (Fig. 1B).

The most differentially expressed genes were those involved in protein synthesis, where 50 genes were upregulated, while 11 were downregulated. Among the genes upregulated, many are related to tRNA synthesis and ribosomal proteins, such as *rpsU*-1, *rpsB*, *rpsB*, *rpsT*, *rpmB*, *rplU* and *rpsL*. The second group corresponds to genes involved in regulatory functions and transcription with 55 genes (42 upregulated and 13 downregulated). The high number of upregulated transcriptional regulators points out the response of this bacterium to a non-common metal as electron acceptor. The third group of genes is involved in energy metabolism and electron transport (12 upregulated and 29 downregulated), of which 20 code for *c*-type cytochromes.

### Expression of *c*-type cytochromes genes during Pd(II) reduction

About half of the differentially expressed genes involved in energy metabolism and electron transport were related to *c*-type cytochromes (9 upregulated and 11 downregulated); 8 are located in the outer membrane, 8 in the periplasmic, 1 in the cytoplasm, and 1 attached to the inner membrane, while the location of the remaining 2 is unknown (Table 2).

**TABLE 2.** *c*-type cytochrome and putative cytochrome genes that were differentially expressed in palladium reduction

**Cytochromes genes that were upregulated in RNA-Seq experiments**
| Locus ID | Gene annotation | Gene | Fold | log2 FC | P-value | Signal | Cellular location |
| --- | --- | --- | --- | --- | --- | --- | --- |
|  |  | name | change |  |  | peptide |  |
| GSU0612 | Cytochromec3, 3 heme-binding sites | *ppcA* | 0.381 | 1.390 | 1.4E-06 | Yes | Periplasmic |
| GSU1024 | Cytochromec3, 3 heme-binding sites | *ppcD* | 0.440 | 1.186 | 2.4E-03 | Yes | Periplasmic |
| GSU1062 | Cytochrome c putative, 1 heme-binding site | - | 0.281 | 1.830 | 4.1E-05 | Yes | Periplasmic |
| GSU2513 | Cytochrome c family protein, 1 heme-binding site | - | 0.423 | 1.243 | 2.4E-02 | Yes | Unknown |
| GSU2808 | Cytochrome c family protein, 5 heme-binding sites | - | 0.338 | 1.563 | 3.0E-05 | Yes | Outer membrane |
| GSU2934 | Cytochrome c family protein, 9 heme-binding sites | *cbcN* | 0.476 | 1.071 | 2.6E-04 | Yes | Periplasmic |
| GSU3107 | Ribosomal protein L31, 1 heme-binding site | *rpmE* | 0.404 | 1.307 | 5.5E-03 | NO | Cytoplasmic |
| GSU2883 | Cytochrome c family protein, 18 heme-binding sites | *omcH<sup>a</sup>* | 0.789 | 0.342 | 1.0E+00 | Yes | Outer membrane |
| GSU2294 | Cytochrome c family protein, 4 heme-binding sites | *omcM<sup>a</sup>* | 0.530 | 0.917 | 5.8E-02 | Yes | Outer membrane |

**Cytochromes genes that were downregulated in RNA-Seq experiments**
| Locus ID | Gene annotation | Gene | Fold | log2 FC | P-value | Signal | Cellular location |
| --- | --- | --- | --- | --- | --- | --- | --- |
|  |  | name | change |  |  | peptide |  |
| GSU0283 | Sensor histidine kinase, 1 heme-binding site | - | 2.442 | -1.288 | 7.9E-05 | NO | Cytoplasmic membrane |
| GSU0592 | Cytochrome c family protein, 11 heme-binding sites | *omcQ* | 2.491 | -1.317 | 1.6E-05 | Yes | Outer membrane |
| GSU0615 | Cytochrome c family protein, 5 heme-binding sites | - | 4.283 | -2.099 | 1.9E-03 | Yes | Periplasmic |
| GSU2076 | Cytochrome c family protein, 5 heme-binding sites | *omcZ* | 2.181 | -1.125 | 4.8E-06 | Yes | Outer membrane |
| GSU2495 | Cytochrome c family protein, 22 heme-binding sites | - | 3.333 | -1.737 | 2.5E-03 | NO | Unknown |
| GSU2731 | Polyheme membrane-associated cytochrome c, 8 heme-binding sites | *omcC* | 2.125 | -1.088 | 2.3E-03 | Yes | Outer membrane |
| GSU2737 | Polyheme membrane-associated cytochrome c, 8 heme-binding sites | *omcB* | 4.047 | -2.017 | 4.6E-02 | Yes | Outer membrane |
| GSU2811 | Cytochrome c Hsc, 1 heme-binding site | *cccA* | 2.657 | -1.410 | 6.8E-16 | Yes | Periplasmic |
| GSU2813 | Cytochrome c551 peroxidase, 2 heme-binding sites | *ccpA* | 3.285 | -1.716 | 4.5E-13 | Yes | Periplasmic |
| GSU2937 | Cytochrome c family protein, 5 heme-binding sites | - | 3.523 | -1.817 | 9.9E-03 | Yes | Periplasmic |
| GSU2504 | Cytochrome c family protein, 1 heme-binding site | *omcS*<sup>a</sup> | 1.829 | -0.871 | 1.0E+00 | Yes | Outer membrane |

Protein cellular location was predicted with PSORT-B (49) and SignalP 3.0 Server (50).
<sup>a</sup> values are out of cut off for Fold change but validated by RT-qPCR

The most upregulated *c*-type cytochromes under Pd(II)-reducing conditions were GSU1062, GSU2808 and PpcA. GSU1062, a putative *c*-type cytochrome, is abundant under ferric citrate reducing conditions (27). Similarly, *gsu2808* encodes for an outer membrane cytochrome that it is reported overexpressed under Fe(III)-reducing conditions and its expression decreases in OmcB deficient conditions (28, 29). On the other hand, cytochrome PpcA participates in electron transfer in the periplasm. A *ppcA* mutant strain showed a decrease in Fe(III) and U(VI) reduction (30, 10). Additionally, the outer membrane cytochromes OmcH and OmcM were also overexpressed under Pd(II)-reducing conditions. Previous work has shown that *omcH* was overexpressed under growing conditions with insoluble Fe(III) oxides; while mutations in the *omcH* and *omcM* genes compromise the reduction of Fe(III) oxides (31). Overexpression of these cytochromes under Pd(II)-reducing conditions suggests that they could be involved in the extracellular reduction of Pd(II).

Other *c*-type cytochromes upregulated during Pd(II) reduction were PpcD, GSU2513, GSU2934 and GSU3107. PpcD is overexpressed during the reduction of Mn(IV) oxides, while the *gsu2934* gene is overexpressed during reduction of Fe(III) oxides (31). The putative cytochrome GSU2513 has not previously been reported and its function has yet to be elucidated.

Surprisingly, during Pd(II) reduction, cytochromes OmcB, OmcC, OmcS and OmcZ, which are involved in the reduction of Fe(III), Mn(IV), and U(VI) (31) were downregulated. A previous model of microbial reduction of Pd(II) by *G. sulfurreducens* suggested that cytochromes OmcB and OmcS, important in the reduction of Fe(III), Mn(IV) or U(VI), could be involved in the reduction process (8); however, our data suggest that the Pd(II) reduction pathway does not involve those common cytochromes, but others with different biochemical characteristics.

### Transcriptional regulation genes expressed in Palladium reduction

The expression profile of some genes encoding proteins involved in transcriptional regulation was considerably altered under Pd(II)-reducing conditions (Table 3). The *gnfK*, *gnfR* and *glnB* genes were upregulated. It has previously been reported that *gnfK*, *gnrR* and *glnB* are upregulated during nitrogen fixation in *G. sulfurreducens* (32). *gnfK* and *gnfR* encode for a un orthodox two-component system, being GnfK a Histidine Kinase and GnrR a Response Regulator. Phosphorylated GnfR, instead of acting as a transcriptional regulator, as the majority of the Response Regulators, binds to the *nifH* mRNA, preventing the formation of a stem-loop structure and therefore avoiding transcription termination (32). In *Geobacter* species, overexpression of genes related to nitrogen fixation has been detected in sediments contaminated with crude oil, groundwater contaminated with uranium, and in microbial fuel cells (33, 34). Under such conditions, where nutrient limitation may exist, it has been proposed that nitrogen fixation is critical for cell growth. Similarly, nitrogen fixation may be an important process during Pd(II) reduction.

**TABLE 3.** Genes encoding transcriptional regulators differentially expressed in during palladium reduction

**Genes encoding proteins involved in transcriptional regulation that were upregulated in RNA-Seq experiments**
| **Locus ID** | **Gene annotation** | **Gene name** | **Fold change** | **log**<br>**FC** | **P-value** |
| --- | --- | --- | --- | --- | --- |
| GSU0284 | TraR/DksA family, zinc finger transcriptional regulator | *dksA* | 0.324 | 1.63 | 5.7E-05 |
| GSU0298 | Two-component system | *fgrM-N* | 0.347 | 1.53 | 8.0E-23 |
| GSU0475 | Sensor histidine kinase, PAS domain-containing | - | 0.401 | 1.32 | 1.4E-02 |
| GSU0596 | Response receiver | - | 0.304 | 1.72 | 2.1E-11 |
| GSU0849 | Regulator of cell morphogenesis and NO signaling | *scdA* | 0.328 | 1.61 | 2.8E-03 |
| GSU0863 | Regulatory protein | - | 0.292 | 1.78 | 2.7E-03 |
| GSU0877 | Response regulator, PilZ domain-containing | - | 0.203 | 2.30 | 7.4E-04 |
| GSU0941 | Nitrogen fixation transcript antitermination sensor histidine kinase | *gnfK* | 0.134 | 2.90 | 6.1E-08 |
| GSU0962 | Two-component system, NtrC family, sensor kinase | - | 0.186 | 2.42 | 1.5E-04 |
| GSU1038 | Response receiver histidine kinase | - | 0.338 | 1.57 | 6.2E-03 |
| GSU1072 | IclR family, transcriptional regulator | - | 0.070 | 3.83 | 2.3E-08 |
| GSU1117 | Response regulator | - | 0.243 | 2.04 | 3.5E-04 |
| GSU1345 | BadM/Rrf2 family, transcriptional regulator | - | 0.203 | 2.30 | 2.5E-03 |
| GSU1379 | Transcriptional regulator, ferric uptake regulator | *fur* | 0.275 | 1.86 | 1.0E-07 |
| GSU1419 | Cro/CI family, transcriptional regulator | - | 0.278 | 1.85 | 5.3E-04 |
| GSU1521 | Integration host factor, alpha subunit | *ihfA-1* | 0.250 | 2.00 | 2.8E-14 |
| GSU1639 | BadM/Rrf2 family, transcriptional regulator | - | 0.082 | 3.61 | 1.6E-16 |
| GSU1658 | Response regulator receiver modulated diguanylate cyclase | - | 0.263 | 1.93 | 1.9E-07 |
| GSU1836 | Nitrogen regulatory protein P-II | *glnB* | 0.094 | 3.41 | 1.0E-16 |
| GSU1999 | RNA-binding protein Hfq | *hfq* | 0.481 | 1.06 | 2.5E-03 |
| GSU2016 | Sensor diguanylate cyclase/phosphodiesterase | - | 0.428 | 1.23 | 5.7E-03 |
| GSU2046 | Response regulator | - | 0.436 | 1.20 | 5.0E-05 |
| GSU2287 | response regulator | - | 0.358 | 1.48 | 4.6E-05 |
| GSU2288 | Sensor histidine kinase | - | 0.298 | 1.75 | 1.4E-03 |
| GSU2384 | sensor histidine kinase, GAF domain-containing | - | 0.275 | 1.86 | 2.3E-03 |
| GSU2625 | ArsR family, transcriptional regulator | - | 0.194 | 2.36 | 2.0E-03 |
| GSU2641 | PATAN domain GTPase-activating protein | - | 0.138 | 2.86 | 1.4E-33 |
| GSU2666 | TetR family, transcriptional regulator | - | 0.443 | 1.17 | 1.3E-05 |
| GSU2735 | TetR family, transcriptional regulator | - | 0.268 | 1.90 | 1.8E-03 |
| GSU2741 | TetR family, transcriptional regulator | - | 0.200 | 2.32 | 1.8E-02 |
| GSU2809 | Fur family, transcriptional regulator | - | 0.333 | 1.59 | 1.3E-05 |
| GSU2822 | Response regulator receiver and ANTAR domain protein | *gnfR* | 0.055 | 4.19 | 4.5E-06 |
| GSU2980 | CopG family transcriptional regulator, nickel-responsive regulator | *nikR* | 0.117 | 3.09 | 2.9E-04 |
| GSU3127 | AraC family, transcriptional regulator | - | 0.441 | 1.18 | 5.0E-02 |
| GSU3206 | TraR/DksA family, zinc finger transcriptional regulator | - | 0.147 | 2.77 | 9.0E-07 |
| GSU3261 | Response regulator | - | 0.439 | 1.19 | 2.9E-04 |
| GSU3292 | Fur family, transcriptional regulator | - | 0.438 | 1.19 | 1.9E-07 |
| GSU3298 | Cro/CI family, transcriptional regulator | - | 0.208 | 2.26 | 8.4E-04 |
| GSU3364 | Transcriptional regulator | *hgtR* | 0.021 | 5.54 | 3.6E-16 |
| GSU2185 | FlgM family, anti-sigma-28 factor | - | 0.135 | 2.89 | 9.9E-07 |

**Genes encoding proteins involved in transcriptional regulation that were downregulated in RNAseq experiments**
| Locus ID | Gene annotation | Gene name | Fold change | log FC | P-value |
| --- | --- | --- | --- | --- | --- |
| GSU0013 | MarR family, transcriptional regulator | - | 3.228 | -1.69 | 5.9E-08 |
| GSU0452 | Sensor histidine kinase | - | 2.315 | -1.21 | 4.5E-04 |
| GSU0534 | BadM/Rrf2 family, transcriptional regulator | *iscR-1* | 2.065 | -1.05 | 5.8E-12 |
| GSU0537 | Sensor diguanylate cyclase/phosphodiesterase, PAS domain-containing | - | 2.134 | -1.09 | 1.1E-03 |
| GSU0811 | Fis family, transcriptional regulator | - | 3.363 | -1.75 | 4.1E-02 |
| GSU0836 | RNA polymerase-binding protein Rnk | *rnk-2* | 91.515 | -6.52 | 4.2E-28 |
| GSU0837 | Response regulator | - | 99.620 | -6.64 | 2.0E-30 |
| GSU0841 | NtrC family, response regulator | - | 39.998 | -5.32 | 4.3E-04 |
| GSU0842 | Sensor histidine kinase response regulator, GAF and PAS domain-containing | - | 15.129 | -3.92 | 3.6E-46 |
| GSU1119 | Response receiver histidine kinase | - | 2.417 | -1.27 | 4.7E-03 |
| GSU1671 | Response receiver-modulated diguanylate cyclase | - | 2.223 | -1.15 | 5.6E-04 |
| GSU2602 | Integration host factor, subunit beta | *ihfB-2* | 3.979 | -1.99 | 1.1E-02 |
| GSU2815 | Sensor histidine kinase, PAS domain-containing | - | 2.419 | -1.27 | 3.3E-02 |

Unexpectedly, HgtR was upregulated more than 5.5 times under Pd(II)-reducing conditions. HgtR has been reported as a global regulator for genes involved in biosynthesis and energy generation in *Geobacter* species. Its expression was essential for growth with hydrogen, during which hgtR expression was induced (35). Moreover, it represses the transcription of several genes of the central metabolism and energy generation, such as *gltA* (citrate synthase), *nouA* (NADH dehydrogenase I subunit), *atpG* (ATP synthase FoB ‘), *srfB* Reductase B’ subunit), *icd* (isocitrate dehydrogenase) and *gntR* (35).

We observed a decrease in the expression of *gltA*, *icd*-*mdh*, *atpG* and *nuo* genes, likely by the upregulation of *hgtR* under Pd(II)-reducing conditions. The increase in *hgtR* expression in our experiments was presumably due to the presence of Pd(II), and not to hydrogen, because we used strict anaerobic conditions under a N_2_/CO_2_ (80/20%) atmosphere without H_2_. In addition, *nuoH*-1, *nouD*, *nuoL*-1 and *nuoF*-1, which form part of an operon of 14 genes that encode for the NADH dehydrogenase, were also downregulated (see Table S1).

The transcriptional regulator Fur was also upregulated during Pd(II) reduction. Fur activity is controlled by intracellular levels of Fe(II) and directly regulates the expression of the *feoAB*, *gsu2432*, *gsu2937*, *gsu3274*, *ideR* and *gsu1639* genes (36). However, the regulon Fur also represses genes, such as citrate synthase (*gltA*), isocitrate dehydrogenase (*icd*), 2-oxoglutarate oxidoreductase (*gsu1467*, *gsu1468*, *gsu1469* and *gsu1470*), malate dehydrogenase (*mdh*), NADH (*gsu0346*, *gsu0347*, *gsu0348*, *gsu0349*, *gsu0350*, *gsu0351*) and OmcZ (*gsu2076*) (36). Fur upregulation during Pd(II) reduction may be the result of iron limiting condition, so that cells would have been trying to overexpress genes related to homeostasis and iron uptake. With these results, we can observe that during Pd(II) reduction, *gltA*, *icd*, *mdh*, *gsu1466*, *gsu1468*, *gsu0349* and *omcZ* were downregulated and could be the result of Fur and HgtR overexpression.

Among the genes coding for possible transcriptional regulators that are upregulated during Pd(II) reduction are *gsu1072*, *gsu0922*, *gsu1639*, *gsu2980*, *gsu3206*, *gsu2185* whose function has not yet been elucidated. Overall, other genes coding for Histidine Kinases and Response Regulators, as well as transcriptional activators and repressors associated with Pd(II) reduction, were differentially expressed.

### Cytochrome *c* and PilA proteins content during palladium reduction

In order to evaluate if mRNA expression correlates with protein content for some *c*-type cytochromes differentially expressed under Pd(II)-reducing conditions, we examined their level by heme-staining of protein gels. As shown in Figure 2A, preparations of soluble fraction, inner membrane and outer membrane proteins, revealed differences in abundance of *c*-type cytochromes in acetate-Pd(II) vs acetate-fumarate conditions. Those bands corresponded to the OmcB and OmcC outer membrane multiheme *c*-type cytochromes, required for Fe(III) reduction in *G. sulfurreducens* (29, 37). OmcS, which is required for Fe(III) oxide reduction (38) and the outer membrane multiheme *c*-type cytochrome, OmcZ, required for optimal current production in microbial fuel cells (39), were abundant in acetate-fumarate in contrast to acetate-Pd(II) conditions.

**FIG 2.**
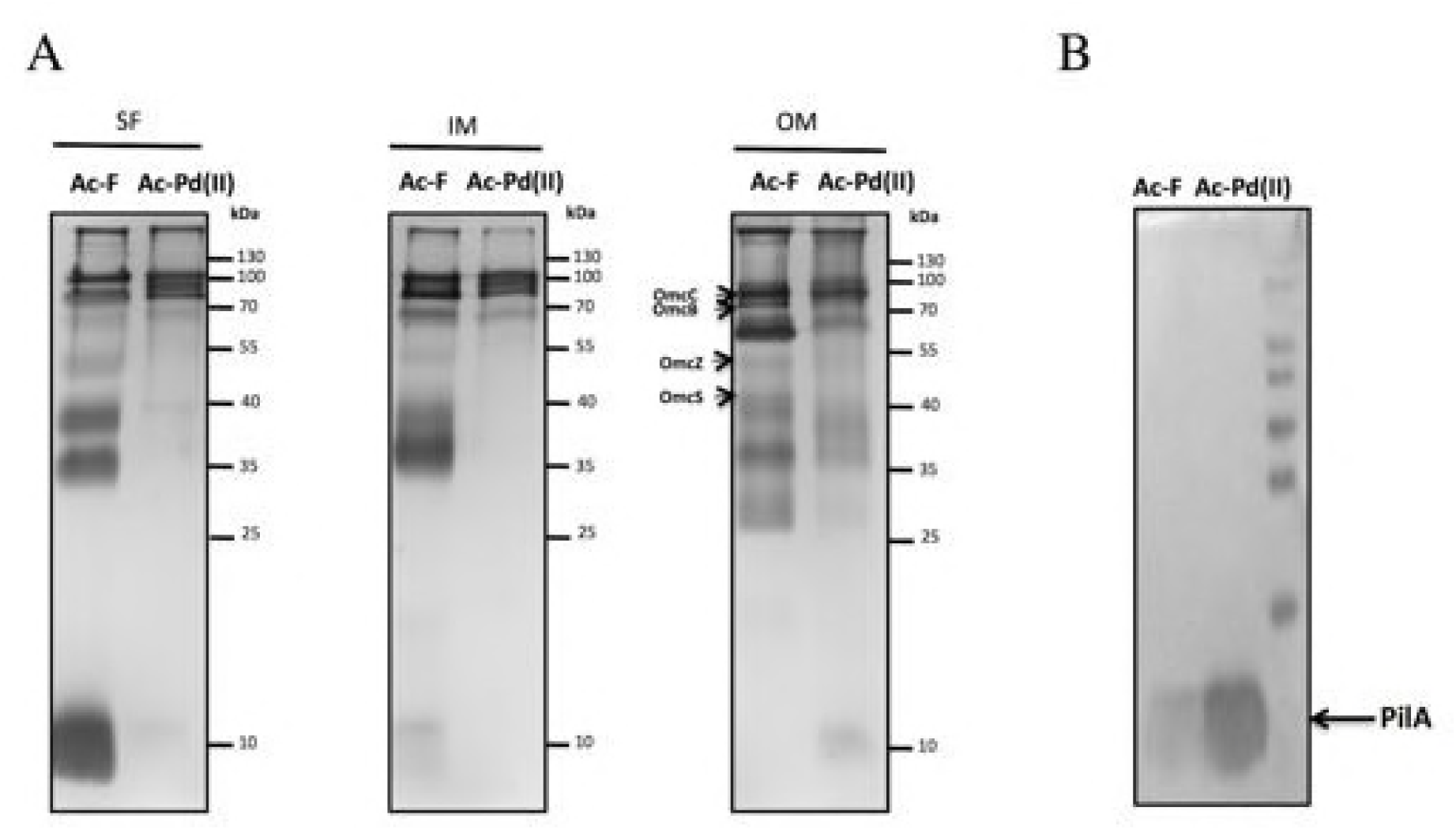
*c*-type cytochrome and PilA protein content found under Pd(II)-reducing conditions. (A) SDS-PAGE heme stained. Outer membrane (OM), inner membrane (IM) and soluble fraction (SF) were prepared from PCA strain with acetate-fumarate (Ac-F) or acetate-Pd(II) (Ac-Pd(II)). The localization of OmcC, OmcB, OmcS, OmcZ and PpcA were labeled based on expected molecular weight (78.96, 74.89, 42.94, 47.09 and 7.72 kDa, respectively). (B) Immunoblot analysis for PilA. The PageRuler Prestained Protein Ladder standard (ThermoScientific) was used as a molecular weight.

We also examined the expression of *pilA* gene (GSU1496), which was upregulated under Pd(II)-reducing conditions. To verify the PilA protein content under these conditions, Immunobloting analysis was performed using anti-PilA antibodies. As shown in Figure 2B, PilA was overproduced under these conditions. These results were surprising since the pili is reported to be required for extracellular electron transfer to insoluble electron acceptors, such as metal oxides and electrodes (12), but not to soluble metals.

### Contribution of pili on Pd(II) reduction

In *G. sulfurreducens*, the pili is an important structure participating in long-distance extracellular electron transfer to Fe(III) oxides, electron exchange between syntrophic partners, as well as electrodes to generate bioelectricity (12, 40, 41). In order to evaluate the contribution of pili to the reduction of Pd(II) to Pd(0), we studied this process in several mutants that have a direct effect on PilA synthesis. Strain Δ*pilR* does not produce the PilR, which is the major transcriptional activator of the *pilA* gene, which codes for pilin, the structural protein of pili (12); therefore, in this mutant, PilA is severely diminished (23). Thus, Pd(II) reduction in Δ*pilA* and Δ*pilR* mutant strains was quantified. We observed that 98% of Pd(II) is reduced to Pd(0) in the WT strain within the first hour of incubation, while in the Δ*pilA* strain, 81% of Pd(II) was reduced during the same incubation period. When we tested the Δ*pilR* strain, only 61% of Pd(II) reduction was reached after three hours (Fig. 3A). Pd(II) reduction was concomitant with an evident change in color of the biomass, which turned black (Fig. 3A inset). To analyze the nature of the palladium NP’s produced, samples of cells covered with these metal were analyzed by XRD (Fig. 3B). The results confirmed the formation of Pd(0) NP’s. The pattern of XRD in all samples showed five strong Bragg reflections at 2 < theta > values around 40.11, 46.66, 68.13, 82.11, which correspond to planes (111), (200), (220), and (311) of a face-centered cubic lattice (fcc) (XRD pattern was indexed to ICDD card 89–4897 (fcc palladium syn)). XRD pattern showed that the Pd NPs were crystalline in nature.

**FIG 3.**
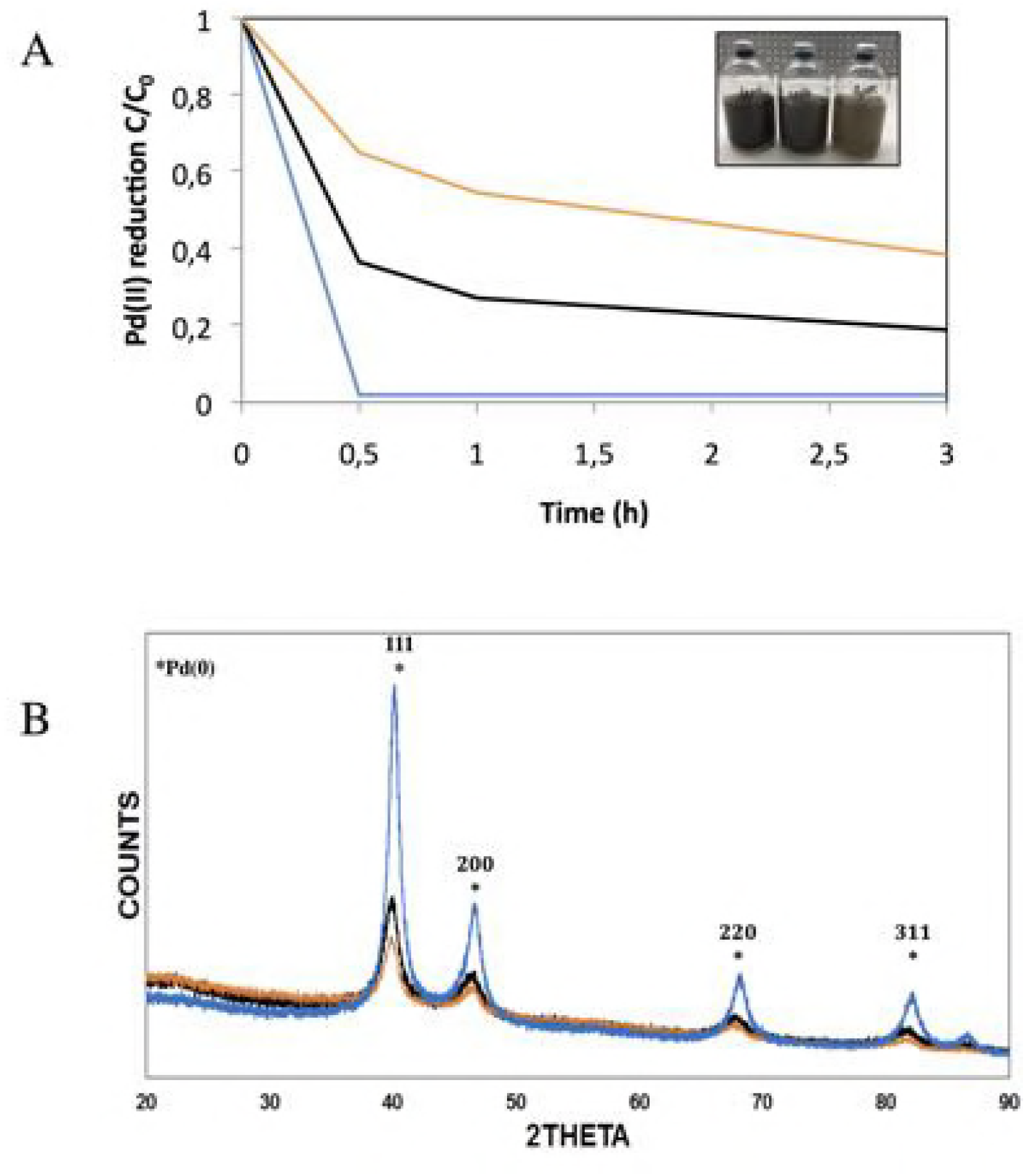
Palladium reduction by different strains of *G. sulfurreducens*. (A) Kinetics of Pd(II) reduction. Photograph of bottles applied for Pd(II) reduction is shown in the inset. (B) Comparison of XRD patters corresponding to cells and black precipitates obtained from cultures of WT, Δ*pilA* and Δ*pilR* strains under Pd(II)-reducing conditions. In (A) and (B), WT strain line blue, Δ*pilA* strain line black, Δ*pilR* strain line orange.

It has been observed that under U(VI)-reducing conditions, using a Δ*pilA* mutant, the production of some *c*-type cytochrome is diminished, which results in a slight decrease in the reduction of U(VI) to U(IV) (42). It has been suggested that the slight reduction of U(VI) in the pili-deficient strain is due to the decrease in outer membrane *c*-type cytochromes and not to pili deficiency (43). Therefore, the slight negative effect on the reduction of Pd(II) by the Δ*pilA* strain, in our experiments, could be due to a decrease in the production of outer membrane *c*-type cytochromes, rather than a negative effect on the pili itself, similarly to reports on U(VI) reduction (43). Since PilR is a transcriptional regulator that controls the expression of at least 44 genes, among which are several *c*-type cytochromes, we propose that the decreased in Pd(II) reduction observed in the Δ*pilR* mutant strain could be related to *c*-type cytochrome content rather than to pili absence. Pd(II) is a toxic metal for many microorganisms; it may inhibit the activity of creatine kinase, aldolase, succinate dehydrogenase, carbonic anhydrase, alkaline phosphatase and prolyl hydroxylase (44). Pd(II) has been found to be toxic in *Shewanella oneidensis*; however, exposed cells could be recovered when a suitable electron donor was provided (1). Similar to *S. oneidensis*, *G. sulfurreducens* could recover viability after the reduction of Pd(II) in NBAF medium, as shown in supplementary data (Fig. S1). In addition, the Δ*pilA* and Δ*pilR* mutant strains took slightly more time than the WT strain to recover viability after exposure to Pd(II) (Fig. S1), which may be due to increased exposure to Pd(II), resulting from the decreased capacity of these strains to reduce this metal (Fig. 3A).

### Growth of *Geobacter sulfurreducens* coupled to Pd(II) reduction

Because the biological reduction of Pd(II) by *S. oneidensis*, *D. desulfuricans* and *G. sulfurreducens* was carried out with resting cells, it is unknown whether this process can be coupled to microbial growth (3, 1, 2). In order to evaluate if *G. sulfurreducens* can couple growth to Pd(II) reduction, incubations were performed in FW medium (see experimental procedures) using acetate (20 mM), lactate (15 mM) or pyruvate (15 mM) as electron donors, and Pd(II) (5 mg/mL) as electron acceptor. No microbial growth was observed with acetate and lactate, which may be due to the fact that the RNA-Seq results showed that the *gltA* (citrate synthase) and *mdh* (malate dehydrogenase) genes were downregulated during Pd(II) reduction, possibly exerting a negative effect on the TCA cycle and therefore on the production of energy. However, by providing pyruvate as an electron donor, we would expect that *G. sulfurreducens* could use an alternative pathway to that of acetate metabolism, redirecting it to biomass biosynthesis (45). However, under these conditions, neither growth nor Pd(II) reduction was observed (data not shown). To verify if the lack of microbial growth observed, could be due to a limitation in the nitrogen source, because it is worth noting that, under Pd(II)-reducing conditions, we observed the overexpression of genes related to nitrogen fixation (*gnfK*, *gnfR* and *glnB*), glutamine was also added to the culture medium. Interestingly, under these conditions, we observed microbial growth in pyruvate-Pd(II) medium (Fig. 4A). Moreover, this growth was coupled to the reduction of 90% Pd(II) to Pd(0) (Fig. 4B) and to the consumption of up to 3 mM of pyruvate (Fig. 4C). Our results imply complex linkages between central metabolism and Pd(II) reduction in *G. sulfurreducens.* Future research is necessary to identify which nutritional elements are conditioning factors to favor the bacterial growth in the presence of other rare metals. However, to our knowledge, this paper reports, for the first time, the microbial reduction of Pd(II) coupled to growth.

**FIG 4.**
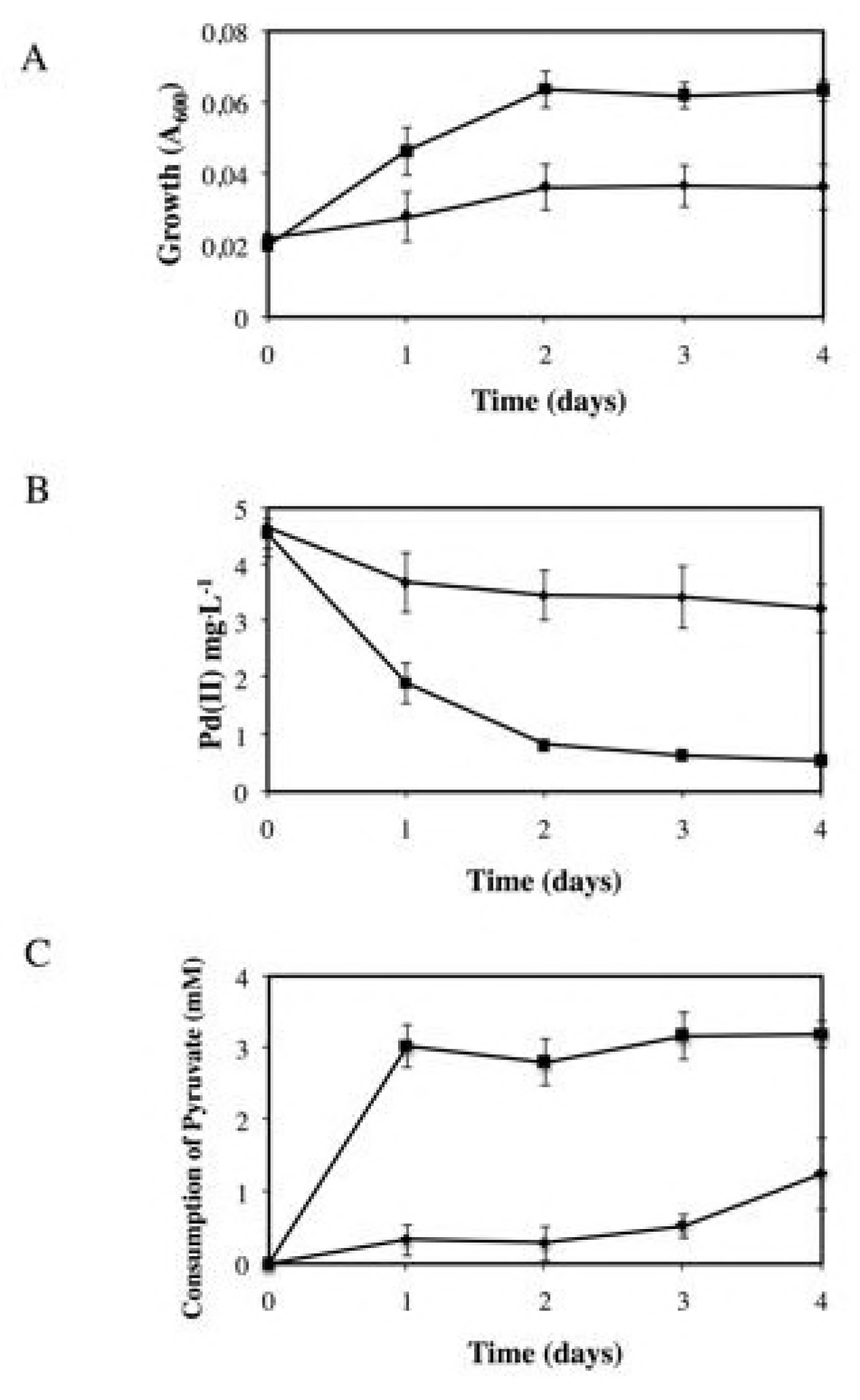
Growth of *Geobacter sulfurreducens* coupled to Pd(II) reduction. (A) Growth. (B) Pd(II) reduction. (C) Consumption of pyruvate. In A, B and C, line with squares, pyruvate-Pd(II) supplemented with glutamine; line with rhombi, pyruvate-Pd(II).

### Validation of selected differentially expressed genes using RT-qPCR

To verify results obtained from RNA-Seq experiments and to get quantitative data to compare the transcript abundances under Pd(II)-reducing conditions, we performed RT-qPCR analyses from 24 selected genes encoding proteins involved in electron transfer, transcriptional regulators and central metabolism (Table 4). These genes included the *c*-type cytochromes genes (*omcH*, *omcM*, *omcB*, *omcC*, *omcS*, *omcZ*, *ppcD*, *gsu1062*, *ppcB*, *omcE*, *ccpA*, *gsu0615*, *gsu2937*, *gsu2513, gsu2808*, *omcQ* and *gsu2495*) and the cold shock DNA/RNA-binding protein, *gsu0207*, the transcriptional regulators, *hgtR* and *gsu0837*, the menaquinol oxidoreductase complex Cbc3, *gsu1650*, the pilin protein, *pilA*, the NADH dehydrogenase I, *nuoH*-1, and the citrate synthase I, *gltA*.

**TABLE 4.** Expression of genes with relevant phenotype observed in RNA-Seq analysis validated by RT-qPCR

| Locus ID | Common name | Avg +Pd/Avg -Pd |
| --- | --- | --- |
| GSU2294 | OmcH, cytochrome c | 2.87 |
| GSU2883 | OmcM, cytochrome c | 1.29 |
| GSU1024 | PpcD, cytochrome c | 5.80 |
| GSU1062 | Cytochrome c, 1 heme-binding site | 5.63 |
| GSU0207 | Cold shock DNA/RNA-binding protein | 12.91 |
| GSU1650 | Cytochrome b/b6 complex | 2.14 |
| GSU3364 | HgtR, hydrogen-dependent growth transcriptional regulator | 9.42 |
| GSU1945 | PilA, pilin protein | 1.50 |
| GSU2513 | Cytochrome c, 1 heme-binding site | 1.91 |
| GSU2808 | Cytochrome c, 5 heme-binding site | 9.44 |
| GSU2737 | OmcB, lipoprotein cytochrome c | 0.28 |
| GSU3731 | OmcC, lipoprotein cytochrome c | 0.33 |
| GSU2076 | OmcZ, cytochrome c | 0.28 |
| GSU0618 | OmcE, cytochrome c | 0.88 |
| GSU2504 | OmcS, cytochrome c | 0.56 |
| GSU0364 | PpcB, cytochrome c | 0.98 |
| GSU0592 | OmcQ, cytochrome c | 0.40 |
| GSU2495 | Cytochrome c, 22 heme-binding site | 0.20 |
| GSU0837 | Response regulator | 0.04 |
| GSU0345 | nuoH-1, NADH dehydrogenase I | 0.74 |
| GSU2813 | CcpA, cytochrome c peroxidase | 0.35 |
| GSU0615 | Cytochrome c, 5 heme-binding site | 0.28 |
| GSU2937 | Cytochrome c, 5 heme-binding site | 0.49 |
| GSU1106 | GltA, type I citrate synthase | 0.48 |

n-fold change were calculated based on the $2^{-DDCT}$ method (21). Avg= Average; -Pd= without palladium and +Pd= with palladium.

Upregulation of *omcH*, *omcM*, *gsu1062*, *gsu2513*, *gsu2808* and *ppcD* genes that encoded for *c*-type cytochromes under Pd(II)-reducing conditions was confirmed by RT-qPCR. Similarly, the expression of *pilA*, *gsu0207*, *hgtR* y *gsu1650* was high under these conditions according to the RT-qPCR. On the other hand, the low transcription of *omcB*, *omcC*, *omcS*, *omcZ*, *gsu0837*, *gsu0345*, *gsu2813*, *gsu0615*, *gsu2937*, *omcQ*, *gsu2495* and *gsu1106* observed in RNA-Seq analyses was confirmed by RT-qPCR. Furthermore, the low transcription of *nuoH*-1 and *gltA* genes was also observed, in agreement with the high expression of *hgtR*, a negative regulator for these genes (35).

### Model for biological Pd(II) reduction by *Geobacter sulfurreducens*

Biological reduction of Pd(II) is a poorly studied process. To date, several bacteria have been reported with the ability to reduce Pd(II) to Pd(0) NP’s (46). In *Desulfovibrio fructosivorans* and *Escherichia coli*, the biological reduction of Pd(II) to Pd(0) is linked to the activity of a hydrogenase (47, 48). *Shewanella oneidensis* and *D. desulfuricans* present a similar Pd(II) reduction mechanism, suggesting that hydrogenase and cytochrome c3 are involved in the reduction process (3, 46).

Based on the results shown here, we propose the following Pd(II) reduction model: Electrons derived from the anaerobic metabolism are transferred through the electron carriers NADH dehydrogenase and MQ to the periplasmic cytochrome MacA, which transfers the electrons to the tri-heme-periplasmic cytochromes PpcA/PpcD. PpcA/PpcD then transport the electrons to the outer membrane cytochromes GSU2513, GSU2808, OmcM and OmcH, which ultimately reduce Pd(II) to Pd(0) (Fig. 5). In addition, the periplasmic cytochromes GSU1062 and CbcN could be intermediates in electron transfer from MacA and/or PpcA/PpcD to the outer membrane cytochromes. Future analyses are necessary to elucidate whether Pd(II) reduction proceeds in the periplasm.

**FIG 5.**
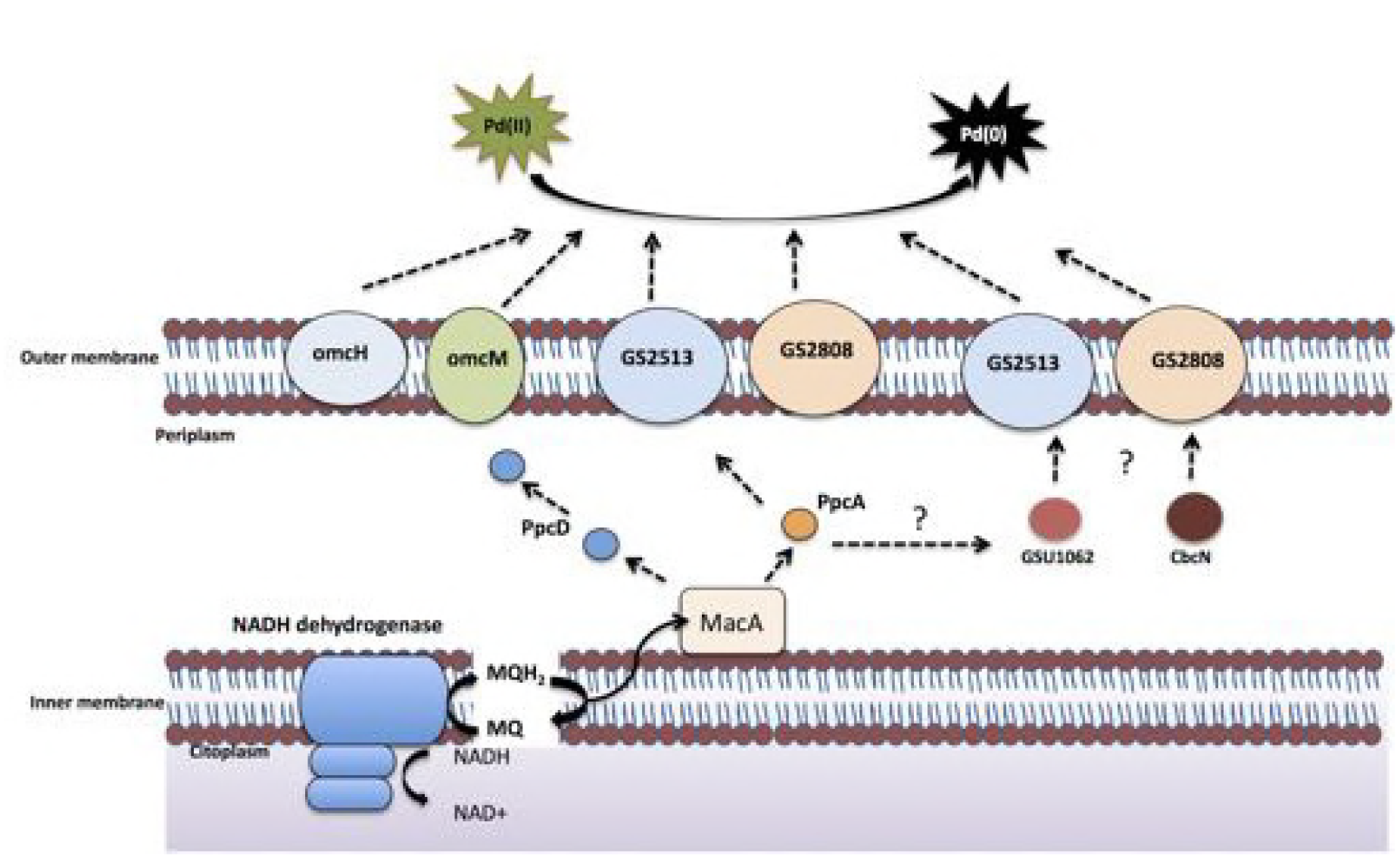
Model of Pd(II) reduction by *Geobacter sulfurreducens*.

## Acknowledgments

We thank Raunel Tinoco, Ramiro Baeza Jimenez, Ricardo Grande and Veronica Jiménez for their technical support. This study was financially supported by CONACYT (Program Frontiers in Science, Grant 1289 and Program Basic Science, Grant 255476). We thank Enrique Morett for critically reading the manuscript. Finally, we greatly acknowledge the support from the national laboratories USMB, LANBAMA and LINAN for their contribution in sample analyses.

## References

1. De Windt W, Boon N, Van den Bulcke J, Rubberecht L, Prata F, Mast J, Hennebel T, Verstraete W. 2006. Biological control of the size and reactivity of catalytic Pd(0) produced by *Shewanella oneidensis*. Antonie van Leeuwenhoek 90:377–389.

2. Pat-Espadas AM, Razo-Flores E, Rangel-Mendez JR, Cervantes FJ. 2013. Reduction of palladium and production of nanocatalyst by *Geobacter sulfurreducens*. Appl Microbiol Biotech 97:9553–9560.

3. Lloyd JR, Yong P, Macaskie LE. 1998. Enzymatic recovery of elemental palladium by using sulfate-reducing bacteria. Appl Environ Microbiol 64(11):4607–4609.

4. Caccavo F, Lonergan DJ, Lovley DR, Davis M, Stolz JF, McInerney MJ. 1994. *Geobacter sulfurreducens* sp. nov., a hydrogen-and acetate-oxidizing dissimilatory metal-reducing microorganism. Appl Environ Microbiol 60:3752–3759.

5. Sanford RA, Wu Q, Sung Y, Thomas SH, Amos BK, Prince EK, Löffler FE. 2007. Hexavalent uranium supports growth of *Anaeromyxobacter dehalogenans* and *Geobacter* spp. with lower than predicted biomass yields. Environ Microbiol 9(11):2885–2893.

6. Law N, Ansari S, Livens FR, Renshaw JC, Lloyd JR. 2008. Formation of nanoscale elemental silver particles via enzymatic reduction by *Geobacter sulfurreducens*. Appl Environ Microbiol 74(22):7090–7093.

7. Lovley DR, Ueki T, Zhang T, Malvankar NS, Shrestha PM, Flanagan KA, Aklujkar M, Butler JE, Giloteaux L, Rotaru AE, Holmes DE, Franks AE, Orellana R, Risso C, Nevin K P. 2011. Geobacter: the microbe electric’s physiology, ecology, and practical applications. Adv Microb Physiol 59:1–100.

8. Pat-Espadas AM, Razo-Flores E, Rangel-Mendez JR, Cervantes FJ. 2014. Direct and Quinone-Mediated Palladium Reduction by *Geobacter sulfurreducens*: Mechanisms and Modeling. Environ Sci Technol 48(5):2910–2919.

9. Ding YH, Hixson KK, Aklujkar MA, Lipton MS, Smith RD, Lovley DR, Mester T. 2008. Proteome of *Geobacter sulfurreducens* grown with Fe(III) oxide or Fe(III) citrate as the electron acceptor. Biochim Biophys Acta 1784(12): 1935–1941.

10. Mehta T, Coppi MV, Childers SE, Lovley DR. 2005. Outer membrane *c*-type cytochromes required for Fe(III) and Mn(IV) oxide reduction in *Geobacter sulfurreducens*. Appl Environ Microbiol 71(12):8634–8641.

11. Childers SE, Ciufo S, Lovley DR. 2002. *Geobacter metallireducens* accesses insoluble Fe(III) oxide by chemotaxis. Nature 416 (6882):767–769.

12. Reguera G, McCarthy KD, Mehta T, Nicoll JS, Tuominen MT, Lovley DR. 2005. Extracellular electron transfer via microbial nanowires. Nature 435:1098–1101.

13. Shelobolina ES, Coppi MV, Korenevsky AA, DiDonato LN, Sullivan SA, Konishi H, Xu H, Leang C, Butler JE, Kim BC, Lovley DR. 2007. Importance of *c*-type cytochromes for U(VI) reduction by *Geobacter sulfurreducens*. BMC Microbiol 7:16.

14. Shi L, Squier TC, Zachara JM, Fredrickson JK. 2007. Respiration of metal (hydr)oxides by *Shewanella* and *Geobacter* : a key role for multihaem c-type cytochromes. Mol Microbiol 65(1):12–20.

15. Coppi MV, Leang C, Sandler SJ, Lovley DR. 2001. Development of a genetic system for *Geobacter sulfurreducens*. Appl Environ Microbiol 67(7):3180–3187.

16. Robinson MD, McCarthy DJ, Smyth GK. 2010. EDGER: a Bioconductor package for differential expression analysis of digital gene expression data. Bioinformatics 26:139–140.

17. Anders S, Huber W. 2010. Differential expression analysis for sequence count data. Genome Biol 11(10):R106.

18. Tarzona S, García-Alcalde F, Dopazo J, Ferrer A, Conesa A. 2011. Differential expression in RNA-Seq: a matter of depth. Genome Res 21(12): 2213–2223.

19. Robinson MD, Oshlack A. 2010. A scaling normalization method for differential expression analysis of RNA-Seq data. Genome Biol 11(3): R25.

20. Kanehisa M, Goto S. 2000. KEGG: Kyoto Encyclopedia of genes and genomes. Nucleic Acids Res 28(1):27–30.

21. Livak K, Shmittgen TD. 2001. Analysis of relative gene expression data using real-time quantitative PCR and the 2-(Delta Delta C(T)) method. Methods 25:402–408.

22. Kim BC, Leang C, Ding YH, Glaven RH, Coppi MV, Lovley DR. 2005. OmcF, a putative *c*-type monoheme outer membrane cytochrome required for the expression of other outer membrane cytochromes in *Geobacter sulfurreducens*. J Bacteriol 187(13):4505–4513.

23. Juárez K, Kim BC, Nevin K, Olvera L, Reguera G, Lovley DR, Methé BA. 2009. PilR, a transcriptional regulator for pilin and other genes required for Fe(III) reduction in *Geobacter sulfurreducens*. J Mol Microbiol Biotechnol 16:146–158.

24. Thomas PE, Ryan D, Levin W. 1976. An improved staining procedures for the detection of the peroxidase activity of cytochrome P-450 on sodium dodecyl sulfate polyacrylamide gels. Anal Biochem 75:168–176.

25. Francis RT Jr, Becker RR. 1984. Specific indication of hemoproteins in polyacrylamide gel using a double-staining process. Anal Biochem 136(2):509–514.

26. Lovley DR, Giovannoni SJ, White DC, Champine JE, Phillips EJP, Gorby YA, Goodwin S. 1993. *Geobacter metallired*ucens gen. nov. sp. nov., a microorganism capable of coupling the complete oxidation of organic compounds to the reduction of iron and other metals. Arch Microbiol 159(4):336–344.

27. Ding YH, Hixson KK, Giometti CS, Stanley A, Esteve-Nuñez A, Khare T, Tollaksen SL, Zhu W, Adkins JN, Lipton MS, Smith RD, Mester T, Lovley DR. 2006. The proteome of dissimilatory metal-reducing microorganism *Geobacter sulfurreducen*s under various growth conditions. Biochim Biophys Acta 1764(7):1198–1206.

28. Methé BA, Webster J, Nevin K, Butler J, Lovley DR. 2005. DNA Microarray analysis of nitrogen fixation and Fe(III) reduction in *Geobacter sulfurreducens*. App Environ Microbiol 71(5): 2530–2538.

29. Leang C, Adams LA, Chin KJ, Nevin KP, Methé BA, Webster J, Sharma ML, Lovley DR. 2005. Adaptation to disruption of the electron transfer pathway for Fe(III) reduction in *Geobacter sulfurreducens*. J Bacteriol 187(17):5918–5926.

30. Lloyd JR, Leang C, Hodges MAL, Coppi MV, Cuifo S, Methe B, Sandler SJ, Lovley DR. 2003. Biochemical and genetic characterization of PpcA, a periplasmic *c*-type cytochrome in *Geobacter sulfurreducens*. Biochem J 369(1):153–161.

31. Aklujkar M, Coppi MV, Leang C, Kim BC, Chavan MA, Perpetua LA, Giloteaux L, Liu A, Holmes DE. 2013. Proteins involved in electron transfer to Fe(III) and Mn(IV) oxides by *Geobacter sulfurreducens* and *Geobacter uraniireducens*. Microbiol 159:515–535.

32. Ueki T, Lovley DR. 2010. Novel regulatory cascades controlling expression of nitrogen-fixation genes in *Geobacter sulfurreducens*. Nucleic Acids Res 38(21):7485–7499.

33. Holmes DE, Nevin KP, Lovley DR. 2004. In situ expression of *nifD* in Geobacteraceae in subsurface sediments. Appl Environ Microbiol 70:7251–7259.

34. Holmes DE, O’Neil RA, Chavan MA, N’Guessan LA, Vrionis HA, Perpetua LA, Larrahondo MJ, DiDonato R, Liu A, Lovley DR. 2009. Transcriptome of *Geobacter uraniireducens* growing in uranium-contaminated subsurface sediments. ISME J 3:216–230.

35. Ueki T, Lovley DR. 2010. Genome-wide gene regulation of biosynthesis and energy generation by a novel transcriptional repressor in Geobacter species. Nucleic Acids Res 38(3):810–821.

36. Embree M, Qiu Y, Shieu W, Nagajaran H, O’Neil R, Lovley DR, Zengler K. 2014. The iron stimulon and Fur regulon of *Geobacter sulfurreducens* and their role in energy metabolism. App Environ Microbiol 80(9):2918–2927.

37. Liu Y, Fredrickson JK, Zachara JM, Shi L. 2015. Direct involvement of ombB, omaB, and omcB genes in extracellular reduction of Fe(III) by *Geobacter sulfurreducens* PCA. Front Microbiol 6:1075.

38. Qian X, Mester T, Morgado L, Arakawa T, Sharma ML, Inoue K, Joseph C, Salgueiro CA, Maroney MJ, Lovley DR. 2011. Biochemical characterization of purified OmcS, a *c*-type cytochrome required for insoluble Fe(III) reduction in *Geobacter sulfurreducens*. Biochim Biophys Acta 1807(4):404–412.

39. Inoue K, Leang C, Franks AE, Woodard TL, Nevin KP, Lovley DR. 2011. Specific localization of the *c*-type cytochrome OmcZ at the anode surface in current-producing biofilms of *Geobacter sulfurreducens*. Environ Microbiol Rep 3(2):211–217.

40. Smith JA, Lovley DR, Tremblay PL. 2013. Outer cell surface components essential for Fe(III) oxide reduction by *Geobacter metallireducens*. Appl Environ Microbiol 79:901–907.

41. Summers ZM, Fogarty HE, Leang C, Franks AE, Malvankar NS, Lovley DR. 2010. Direct exchange of electrons within aggregates of an evolved syntrophic coculture of anaerobic bacteria. Science 330:1413–1415.

42. Cologgi DL, Lampa-Pastirk S, Speers AM, Kelly SD, Reguera G. 2011. Extracellular reduction of uranium via *Geobacter* conductive pili as a protective cellular mechanism. Proc Natl Acad Sci 108(37):15248–15252.

43. Orellana R, Leavitt J, Comolli LR, Csencsits R, Janot N, Flanagan KA, Gray AS, Leang C, Izallalen M, Mester T, Lovley DR. 2013. U(VI) Reduction by diverse outer surface *c*-type cytochromes of *Geobacter sulfurreducens*. Appl Environ Microbiol 79:6369–6374.

44. Liu TZ, Lee SD, Bhatnagar RS. 1979. Toxicity of palladium. Toxicol Lett 4(6):469–473.

45. Wilkins MJ, Wrighton KC, Nicora CD, Williams KH, McCue LA, Handley KM, Miller C S, Giloteaux L, Montgomery AP, Lovley DR, Banfield JF, Long PE, Lipton MS. 2013. Fluctuations in species-level protein expression occur during element and nutrient cycling in the subsurface. PLoS One 8(3): e57819

46. De Corte S, Hennebel T, De Gusseme B, Verstraete W, Boon N. 2012. Bio-palladium: from metal recovery to catalytic applications. Microb Biotechnol 5(1):5–17.

47. Mikheenko IP, Rousset M, Dementin S, Macaskie LE. 2008. Bioaccumulation of palladium by *Desulfovibrio fructosivorans* wild-type and hydrogenase-deficient strains. Appl Environ Microbiol 74: 6144–6146.

48. Deplanche K, Caldelari I, Mikheenko IP, Sargent F, Macaskie LE. 2010. Involvement of hydrogenases in the formation of highly catalytic Pd(0) nanoparticles by bioreduction of Pd(II) using *Escherichia coli* mutant strains. Microbiol 156(9):2630–2640.

49. Gardy JL, Spencer C, Wang K, Ester M, Tusnády GE, Simon I, Hua S, deFays K, Lambert C, Nakai K, Brinkman FS. 2003. PSORT-B: Improving protein subcellular localization prediction for Gram-negative bacteria. Nucleic Acids Res 31(13):3613–3617.

50. Emanuelsson O, Brunak S, von Heijne G, Nielsen H. 2007. Locating proteins in the cell using TargetP, SignalP, and related tools. Nat Protoc 2(4):953–971.

